# Acclimation Capacity to Global Warming of Amphibians and Freshwater Fishes: Drivers, Patterns, and Data Limitations

**DOI:** 10.1101/2023.12.22.573141

**Authors:** Katharina Ruthsatz, Flemming Dahlke, Katharina Alter, Sylke Wohlrab, Paula C. Eterovick, Mariana L. Lyra, Sven Gippner, Steven J. Cooke, Myron A. Peck

## Abstract

Amphibians and fishes play a central role in shaping the structure and function of freshwater environments. These organisms have a limited capacity to disperse across different habitats and the thermal buffer offered by freshwater systems is small. Understanding determinants and patterns of their physiological sensitivity across life history is, therefore, imperative to predicting the impacts of climate change in freshwater systems. Based on a systematic literature review including 345 experiments with 998 estimates on 96 amphibian (Anura/Caudata) and 93 freshwater fish species (Teleostei), we conducted a quantitative synthesis to explore phylogenetic, ontogenetic, and biogeographic (thermal adaptation) patterns in upper thermal tolerance (CT_max_) and thermal acclimation capacity (Acclimation Response Ratio, ARR) as well as the influence of the methodology used to assess these thermal traits using a conditional inference tree analysis. We found globally consistent patterns in CT_max_ and ARR, with phylogeny (taxa/order), experimental methodology, climatic origin, and life stage as significant determinants of thermal traits. The analysis demonstrated that CT_max_ does not primarily depend on the climatic origin but on experimental acclimation temperature and duration, and life stage. Higher acclimation temperatures and longer acclimation times led to higher CT_max_ values, whereby Anuran larvae revealed a higher CT_max_ than older life stages. The ARR of freshwater fishes was more than twice that of amphibians. Differences in ARR between life stages were not significant. In addition to phylogenetic differences, we found that ARR also depended on acclimation duration, ramping rate, and adaptation to local temperature variability. However, the amount of data on early life stages is too small, methodologically inconsistent, and phylogenetically unbalanced to identify potential life cycle bottlenecks in thermal traits. We therefore propose methods to improve the robustness and comparability of CT_max_/ARR data across species and life stages, which is crucial for the conservation of freshwater biodiversity under climate change.

## 1. Introduction

Global climate change is not only causing an increase in mean air and water temperatures, but also an increased magnitude and frequency of extreme climatic events (Lee et al. 2023). As a result, ectotherms are more likely to experience temperatures beyond their critical thermal maximum (CT_max_) in both terrestrial and aquatic habitats (Duarte et al. 2012; Sunday et al. 2014). This is particularly true for populations already living close to their upper thermal limit. Consequently, the ability to mitigate thermal stress through either migration, evolutionary genetic adaptation or acclimation is crucial for the persistence of species in a changing climate (Franks & Hoffmann 2002; Huey et al. 2012; Seebacher et al. 2015). Given the limited dispersal ability of many species (e.g., freshwater species; Woodward et al. 2010) and rapid pace of global warming (Hoffmann & Sgró 2011), physiological acclimation is arguably the most important mechanism for coping with climate change (Gunderson & Leal 2015). Understanding differences in acclimation capacity of species and identifying global patterns can therefore help to identify climate change risks to biodiversity and develop effective conservation measures (Somero 2010).

As an adaptive response to larger seasonal differences in temperature, thermal tolerance and acclimation capacity of ectothermic species or populations tend to increase with increasing latitude from tropical through temperate climate zones (e.g., Somero 2005; Deutsch et al. 2008; Sunday et al. 2011; Peck et al. 2014; Rohr et al. 2018; Cicchino et al. 2023; but see: Sørensen et al. 2016; Gunderson & Stillman 2015) and from higher to lower elevations (Enriquez-Urzelai et al. 2020; but not: Sunday et al. 2019; Gutiérrez-Pesquera et al. 2022). This biogeographical pattern is consistent with the *climate variability hypothesis* (Janzen 1967; Ghalambor et al. 2006), suggesting that climatic differences across altitudinal and latitudinal gradients lead to corresponding adaptations in thermal physiology (but see: Gutiérrez-Pesquera et al. 2022). Low-latitude species adapted to relatively stable temperature conditions may have a lower acclimation capacity and, therefore, may be more vulnerable to climate change (Tewksbury et al. 2008; Sunday et al. 2014; but see: Bovo et al. 2023). However, there is still little empirical evidence supporting the climate variability hypothesis, possibly due to the limited geographical and phylogenetic coverage of observations, and because of the inconsistency of the methods applied to measure the acclimation capacity of different species and life stages (Gutiérrez-Pesquera et al. 2016; Shah et al. 2017). Moreover, the use of different methods or protocols might impact the estimation of thermal traits (Terblanche et al. 2007; Chown et al. 2009; Rohr et al. 2018; Pottier et al. 2022a; but not: Sunday et al. 2019). For example, acclimation duration (i.e., how long organisms were held at an acclimation temperature before being exposed to the test temperature; Rohr et al. 2018; Ruthsatz et al. 2022a) and ramping protocol (i.e., heating or cooling rate in thermal tolerance trials; Illing et al. 2020; Penman et al. 2023) have been suggested to influence measurements of acclimation capacity, as the underlying physiological processes occur over certain time periods.

In animals with complex life histories, thermal tolerance and acclimation capacity are thought to change during ontogeny according to physiological and morphological reorganizations and concomitant aerobic capacities in relation to oxygen demand (Pörtner 2002; Pörtner & Peck 2010; Ruthsatz et al. 2020a,b, 2022a) as well as energetic costs associated with developmental processes (Ruthsatz et al. 2019). Furthermore, life stages might differ in their ability for behavioral thermoregulation (Navas et al. 2008; Little & Seebacher 2017) resulting in stage-specific adaptations in thermal traits (Huey et al. 1999). Therefore, determining taxon-specific acclimation capacity at different ontogenetic stages should be taken into account when studying climate adaptation of ectothermic species, as it will help identify life cycle bottlenecks and provide robust data on the vulnerability of populations or species to global warming (Bodensteiner et al. 2021; Pottier et al. 2022a; Dahlke et al. 2022). To date, most studies have not distinguished among life stages or have only focused on adults when addressing the *climate variability hypothesis* (e.g., Gunderson & Stillman 2015; Sunday et al. 2011, 2014, 2019; Rohr et al. 2018) and the vulnerability of species to global change (e.g., Calosi et al. 2008; Comte & Olden 2017a; Morley et al. 2019; Molina et al. 2023). Moreover, most studies have pooled several pre- and post-metamorphic life stages (Pottier et al. 2022a; Weaving et al. 2022) thereby risking to overlook a critical thermal bottleneck in the life cycle (Dahlke et al. 2020). The extent to which thermal tolerance and acclimation capacity change during ontogeny is, therefore, not clear for many taxa.

Amphibians and freshwater fishes tend to live in relatively shallow waters (e.g., wetlands, ponds, rivers, lakes) and may, therefore, experience strong seasonal and daily temperature fluctuations (Capon et al. 2021) and climate extremes such as heat waves (IPCC 2021). In addition, both taxa have a limited ability to disperse over larger distances and habitats to avoid unfavorable climatic conditions (Albert et al. 2011; Yu et al. 2013; Campos et al. 2021). Consequently, as a result of local adaptation (Meek et al. 2023), there should be a close correspondence between the capacity for thermal acclimation and the climatic conditions that amphibians and freshwater fishes experience during their life cycle. However, freshwater systems offer a wide range of thermal microhabitats that enable behavioral thermoregulation (Campos et al. 2021), especially for (post-metamorphic) amphibians that can switch between water and land. The potential of behavioral thermoregulation could reduce the need for physiological adaptations (also known as the *Bogert Effect*, (Bogert 1949)) and thus counteract the emergence of geographical patterns in thermal acclimation capacity and/or thermal tolerance. Given the central role that amphibians and freshwater fishes play in shaping the structure and function of these ecosystems (Closs et al. 2016; Hocking & Babbitt 2014), understanding determinants and patterns of their physiological sensitivity is imperative to predicting the impacts of climate change on freshwater systems.

Here, we aimed to define the determinants and patterns of acclimation capacity in upper thermal tolerance in amphibians and freshwater fishes addressing, for the first time, ontogeny dependent variation. To do so, we compiled literature on upper thermal tolerance and collected empirical data for CT_max_ in four amphibian (i.e., larvae, metamorphs, juveniles, and adults) and three freshwater fish (i.e., larvae, juveniles, and adults) life stages acclimated to at least two different temperatures. Next, we calculated the population-specific acclimation capacity, i.e., mean acclimation response ratio (ARR) of upper thermal limits (i.e., the slope of the linear function describing the change in thermal tolerance with a given change in acclimation temperature; e.g., Hutchison 1961; Claussen 1977), and conducted a quantitative synthesis on the acclimation capacity of amphibians and freshwater fishes to test for differences among taxonomic groups, among life stages, and across thermal characteristics of populations, i.e., biogeographic differences based on local thermal adaptation. Further, we investigated how the methodological context (i.e., acclimation duration and temperature, ramping rate) affects estimates of thermal traits. Finally, we summarize methodological concerns, highlight key knowledge gaps, and provide research recommendations for generating reliable and comparable data on the acclimation capacity of ectothermic species and their life stages. This will improve our ability to predict future climate vulnerability of species and populations.

## 2. Materials and Methods

### 2.1 Systematic literature review

We conducted a systematic literature review using ISI Web of Science (ISI WOS, 2021) on 2022/06/30 and did not apply a timespan limit. The following Boolean search string was used to capture articles with experimental studies manipulating acclimation temperatures of amphibians and freshwater fishes at different life stages, and subsequently measured their CT_max_: (“amphibian*” OR “newt*” OR “frog*” OR “toad*” OR “salamander*” OR “freshwater" AND "fish*”) AND (“early” OR “young” OR “life stage*” OR “ontogen*” OR “development*” OR “hatchling*” OR “alevin*” OR “larv*” OR “tadpole*” OR "metamorph*" OR “postmetamorph*” OR “post-metamorph*” OR "postlarva*" OR "post-larva*” OR “fry*” OR “parr*” OR “smolt*” OR “subadult*” OR “sub-adult” OR “juvenile” OR “fingerling*” OR “adult*”) AND (“thermal” OR “temperature” OR “acclimat*” OR “heat” OR “warm*”) AND (“tolerance*” OR “thermal tolerance*” OR “temperature tolerance*” OR “warming tolerance*” OR “heat tolerance*” OR “thermal stress tolerance*” OR “heat stress tolerance*” OR “temperature stress tolerance*” OR “limit*” OR “temperature stress*” OR “thermal limit*” OR "critical temperature*" OR “CT max” OR "critical thermal m*" OR “thermal performance breadth*” OR “thermal breadth*” OR “performance breadth*” OR “thermal range*” OR “thermal window*” OR “thermal tolerance window*” OR “tolerance window*” OR "sensitivity*" OR "thermal sensitivity").

Our search resulted in 11,057 documents (Fig. S1). After removing book chapters, conference contributions, reviews, meeting abstracts, editorial material, preprints, and proceedings articles, 10,740 published peer-reviewed articles remained in our initial database. Additionally, we manually added articles included in the meta-analyses of Claussen (1977), Gunderson and Stillman (2015), Comte & Olden (2017b), Morley et al. (2019) and Dahlke et al. (2020) that met our inclusion criteria (specified below) but were not obtained through the ISI Web of Science search. After an initial subjective evaluation of titles, 3,991 articles potentially containing results matching the objective of the present study were kept and further assessed for eligibility using the abstract. Thirty-four articles were not accessible. We contacted the authors of the original articles to request missing information and heard back from two authors. Finally, a total of 93 articles (34 on amphibians; 59 on freshwater fishes) met our inclusion criteria. Search methods are summarized in a PRISMA flowchart (Fig. S1), and a list of included articles and experimental studies therewithin is available in the figshare data repository under https://figshare.com/s/57775032a0b79c416ef2 (DOI: 10.6084/m9.figshare.24872133; published after acceptance).

### 2.2 Inclusion criteria

Experimental studies were selected based on the following seven inclusion criteria:

(1) Experimental studies were conducted on amphibians (anurans, caudates, or gymnophiones) or freshwater fishes (teleosts).
(2) Animals were acclimated to at least two constant acclimation temperatures under laboratory conditions prior to the CT_max_ measurements. Fluctuating temperature treatments were not considered. Therefore, field experiments were not considered since the acclimation temperature under field conditions is likely to fluctuate.
(3) Articles provided comprehensive information on methodology (acclimation temperatures and duration, ramping rate), phylogeny (species names), sampling location (GPS location), and life stage. If no GPS coordinates were provided but a concrete sampling location was stated (e.g., Central Park, New York City, NY, USA), we searched for the coordinates of the respective location on Google Maps.
(4) Animals were collected from their natural habitat. Data were excluded if measurements were taken from specimens bred artificially to reduce confounding issues associated with artificial selective history (Bennett et al. 2018). Experimental studies were also included if adult animals were collected to immediately reproduce in the lab to obtain larvae.
(5) The critical thermal maximum (CT_max_) was used as a standard measure of heat tolerance (Lutterschmidt & Hutchison 1997a). Experimental studies using other measures such as voluntary thermal maximum, time to death, heat knockdown, or lethal temperatures as well as extrapolations from thermal performance curves were not considered. In aquatic and terrestrial ectotherms such as amphibians and fish, CT_max_ is generally measured as loss of equilibrium (LOE) or loss of rightening response (LRR) following a steady increase in water or air temperature [dynamic method according to Fry (1947)]. In comparison to other endpoints such as the onset of spasms (Lutterschmidt & Hutchison 1997b), measuring CT_max_ as LOE or LRR is a non-lethal, robust method at various body sizes that is repeatable within individuals (Morgan et al. 2018).
(6) At CT_max_ measurements, the animals could be classified into one of four different categories representing the consecutive life stages of amphibians and freshwater fish: (a) larva (pre-metamorphic; amphibians: < Gosner stage 42), (b) metamorph (only for amphibians: Gosner stage 42-46), (c) juvenile (post-metamorphic), and (d) adult (after reaching sexual maturity). Embryos were not included in the present study because the assessment of acute heat tolerance in non-mobile life stages requires different endpoints that may not be directly comparable to the LOE/LRR-based CT_max_ of other life stages (Cowan et al. 2023; Lechner et al. 2023).
(7) Food was provided *ad libitum* during the acclimation time since food deprivation might decrease thermal tolerance and/or acclimation capacity (Lee et al. 2016).

### 2.3 Data extraction

When all inclusion criteria were met, data were collated in a spreadsheet. We extracted mean CT_max_ for all acclimation temperatures resulting in 998 single data points (513 for amphibians; 485 for freshwater fish). Some of the selected articles performed different experimental studies on e.g., different populations of one species, different species, or different life stages. Therefore, all available datasets were included, resulting in 345 experimental studies from 93 articles with 345 paired effect sizes CT_max_ and acclimation capacity. Data presented in the text or tables were directly extracted from the articles. When only raw data were available, mean values were calculated. For articles that presented results in figures instead of tables, Engauge Digitizer 12.1 was used (Mitchell et al. 2021) to extract data from the graphs. In addition to CT_max_ data, information on the methodology (i.e., acclimation temperatures and duration, ramping protocol), as well as on variables representing sampling location as detailed as possible (i.e., GPS coordinates), phylogeny (i.e., scientific classification according to the Linnean classification), and life stage at CT_max_ assessment was extracted. The data extractions were performed by KR, KA and PCE, followed by an accuracy check of the data (KA: freshwater fish sub dataset; KR: amphibian sub datatset; PCE: both sub datasets).

### 2.4 Bioclimatic variables

For each sampling location, 19 bioclimatic metrics, related to temperature and precipitation, and elevation were extracted using the WorldClim 2 database (http://www.worldclim.org/; Fick & Hijmans 2017) for the average of the years 1970–2000. The data were extracted at a spatial resolution of 30 seconds (∼1 km^2^), using package ‘geodata’ (Hijmans 2021) in R (version 4.2.1; R Core Team, 2020). Bioclimatic variables are coded as follows (name of the variable and the unit used as input is included in parenthesis): Annual Mean Temperature (Bio1; °C), Mean Diurnal Range (Bio 2; °C), Isothermality (Bio 3; percentage), actual Temperature Seasonality (Bio 4; standard deviation in °C), Maximum Temperature of Warmest Month (Bio 5; °C), Minimum Temperature of Coldest Month (Bio 6; °C), Annual Temperature Range (Bio 7; °C), Mean Temperature of Wettest Quarter (Bio 8; °C), Mean Temperature of Driest Quarter (Bio 9; °C), Mean Temperature of Warmest Quarter (Bio 10; °C), Mean Temperature of Coldest Quarter (Bio 11; °C), Annual Precipitation (Bio 12; mm), Precipitation of Wettest Month (Bio 13; mm), Precipitation of Driest Month (Bio 14; mm), Precipitation Seasonality (Bio 15; coefficient of variation expressed in percent), Precipitation of Wettest Quarter (Bio 16; mm), Precipitation of Driest Quarter (Bio 17; mm), Precipitation of Warmest Quarter (Bio 18; mm), and Precipitation of Coldest Quarter (Bio 19; mm) (http://www.worldclim.org/data/bioclim.html). These macroclimatic data were used as approximations to identify patterns of local adaptation in amphibians and freshwater fish, as microclimatic data (e.g. site-specific temperatures) were not available in original articles or in the WorldClim database. Following previous studies (Morley et al. 2019; Carilo Filho et al. 2022), mean near-surface air temperature was assumed to reflect the temperature profile of freshwater systems and used to analyze thermal adaptation in both taxa. We are aware that these macroclimatic data as provided by the WorldClim database (representing surface air temperature) present some limitations in reflecting microclimatic data (see Conclusion section for methods to improve future studies). In contrast to marine habitats, the temperature of small or shallow bodies of water might fluctuate with the surface air temperature and animals at the surface of freshwater systems might be further exposed to high temperatures. Therefore, we assume that the average near-surface air temperature is a suitable estimate of the temperature of freshwater systems, thereby reflecting the thermal local adaptation of investigated amphibian and freshwater populations. This is in accordance with previous studies testing the effect of thermal adaptation on the thermal physiology of various taxa (e.g., Gutiérrez-Pesquera et al. 2016; Morley et al. 2019; Carilo Filho et al. 2022; Sinai et al. 2022). Sampling locations were assigned to latitudinal groups based on the absolute latitude (°N/S) and were categorized as either tropical (0–25°), sub-tropical (>25–40°), temperate (>40–53.55°) or polar (>53.55°; Morley et al. 2019).

### 2.5 Effect size calculation: acclimation response ratio (ARR)

A well-established method to measure acclimation capacity in thermal tolerance in ectothermic animals is the calculation of the acclimation response ratio (ARR), i.e., the slope of the linear function describing the change in thermal tolerance with a given change in acclimation temperature (e.g., Hutchison 1961; Claussen 1977; Gunderson & Stillman 2015; Morley et al. 2019). We separately calculated the ARR for CT_max_ within each study using the equation according to Claussen (1977):

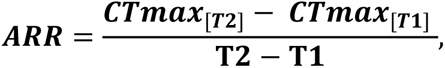

where T represents the acclimation temperature (°C; with T2 = highest acclimation temperature and T1 = lowest acclimation temperature) and CT_max_ the heat tolerance estimates (°C). When data on more than two acclimation temperatures were presented, we calculated the ARR for each stepwise comparison (e.g., 18-20°C, 20-22°C, 22-24°C; Pottier et al. 2022a) and used the mean ARR of all comparisons in the statistical analysis. Higher absolute values of ARR correspond to higher plasticity in thermal tolerance limits (i.e., greater acclimation capacity; Claussen 1977; Kingsolver & Huey 1998; Gunderson & Stillman 2015; van Heerwarden et al. 2016). An acclimation response ratio of 1.00 indicates a 100% acclimation in thermal tolerance to a temperature increase of 1°C (Morley et al. 2019).

### 2.6 Statistical analyses

For all statistical tests R 4.0.2 (R Core Team, 2020) was used. All plots were constructed using R packages ‘ggplot2’ (Wickham & Wickham 2009), ‘ggtree’ (Yu et al. 2017) and Adobe Illustrator CS6.

Conditional inference tree (CIT) analysis (R package ‘partykit’, Hothorn & Zeileis 2015) was used to assess the influence of geographic origin (bioclimatic variables, elevation), phylogeny (taxon and order level), experimental methodology (ramping rate, mean acclimation duration, mean acclimation temperature), and ontogeny on CT_max_ and ARR of amphibians and freshwater fishes. As an advantage over traditional (parametric) methods, CIT is a non-parametric method that handles complex non-linear relationships without making specific assumptions about data distribution or being sensitive to outliers. CIT involves recursive partitioning to split data into subsets based on the relevance of predictor variables. At each node of the tree, a permutation-based test (Monte Carlo method with Bonferroni correction) determines whether the split is statistically significant (α < 0.05). The initial split in the tree indicates which predictor variable has the strongest correlation with the response variable (CT_max_ or ARR). The resulting tree provides a hierarchical structure and classification of significant predictor variables. A post-pruning strategy based on the Akaike Information Criterion (AIC, Akaike 1974) was used to avoid overfitting, i.e., removal of nodes that do not improve the overall fit of the model (Hothorn & Zeileis 2015). The raw data used for the analysis are provided in the electronic supplementary material (Table S1).

Phylogenetic trees for visualizations were created using the R package ‘fishtree’ (Chang et al. 2019) for freshwater fishes (Teleostei). The amphibian ultrametric tree was obtained from the timetree.org website in June 2022 (Kumar et al. 2022). To ensure the validity, we developed a workflow to filter out taxonomically invalid taxa. Firstly, a time tree was generated using the "Build a Timetree" function on timetree.org. The taxa names were then extracted from the generated time trees using the R package ‘ape’ (Paradis & Schliep 2019), excluding non-binomial names. The extracted list was cross-checked with the GBIF Backbone Taxonomy (GBIF Secretariat, 2021) using the species matching tool (https://www.gbif.org/tools/species-lookup, accessed June 2022). Matches at the species rank with an accuracy of 100% were extracted and matches with lower accuracy were manually verified. The resulting species list was uploaded back onto timetree.org using the "Load a List of Species" function. The newly generated time tree was then downloaded and used for visualizations.

The relationship between absolute latitude (°N/S), elevation (m), and the bioclimatic variables that revealed a significant effect on CT_max_ (Bio14, Bio7) and ARR (Bio3) in the CIT analysis was assessed using linear regressions. Differences in respective bioclimatic variables between latitudinal groups (tropical, sub-tropical, temperate, polar) were compared using Kruskal-Wallis tests followed by pairwise Mann-Whitney U-tests with false discovery rate (FDR)-correction.

We were unable to test for a publication bias (i.e., whether statistically insignificant or negative results are less likely to be published) in our data set given the use of a non-standardized effect size metric (i.e., ARR). Therefore, it remains possible that this bias may also occur.

## 3. Results

The analysis of paired CT_max_ and ARR values included 201 experimental studies on 96 amphibian species (2 orders) (Fig. 1a, b) and 144 experimental studies on 93 teleost species (14 orders) (Fig. 1c, d), corresponding to a phylogenetic coverage of approximately 1% within each taxon. No data were available for Gymnophiona. Most experimental studies determined CT_max_ and ARR in adult animals. For freshwater fishes, only one study determining CT_max_ in larvae was obtained, while 20% of the data covered juveniles (Fig. 1c). For amphibian species, 12% of the data was available for one or several of the early-life stages, i.e. larvae, metamorphs, and/or juveniles (Fig. 1a).The geographic origin of amphibian and fish species displayed a bias towards temperate and subtropical regions, with 60% of all sampling sites located in North America, 10% in Australia and New Zealand, 8% in South America, 6% in Europe, and 4% in China (Fig. 1b, d). The CT_max_ protocols were variable in both taxa, with acclimation times ranging from 0.5 to 150 d (mean = 11.2 d), acclimation temperatures ranging from 8.0 to 31.75 °C (mean = 19.45 °C), and ramping rates ranging from 0.02 to 1.0 °C min^-1^ (mean = 0.75 °C min^-1^).

**Figure 1.**
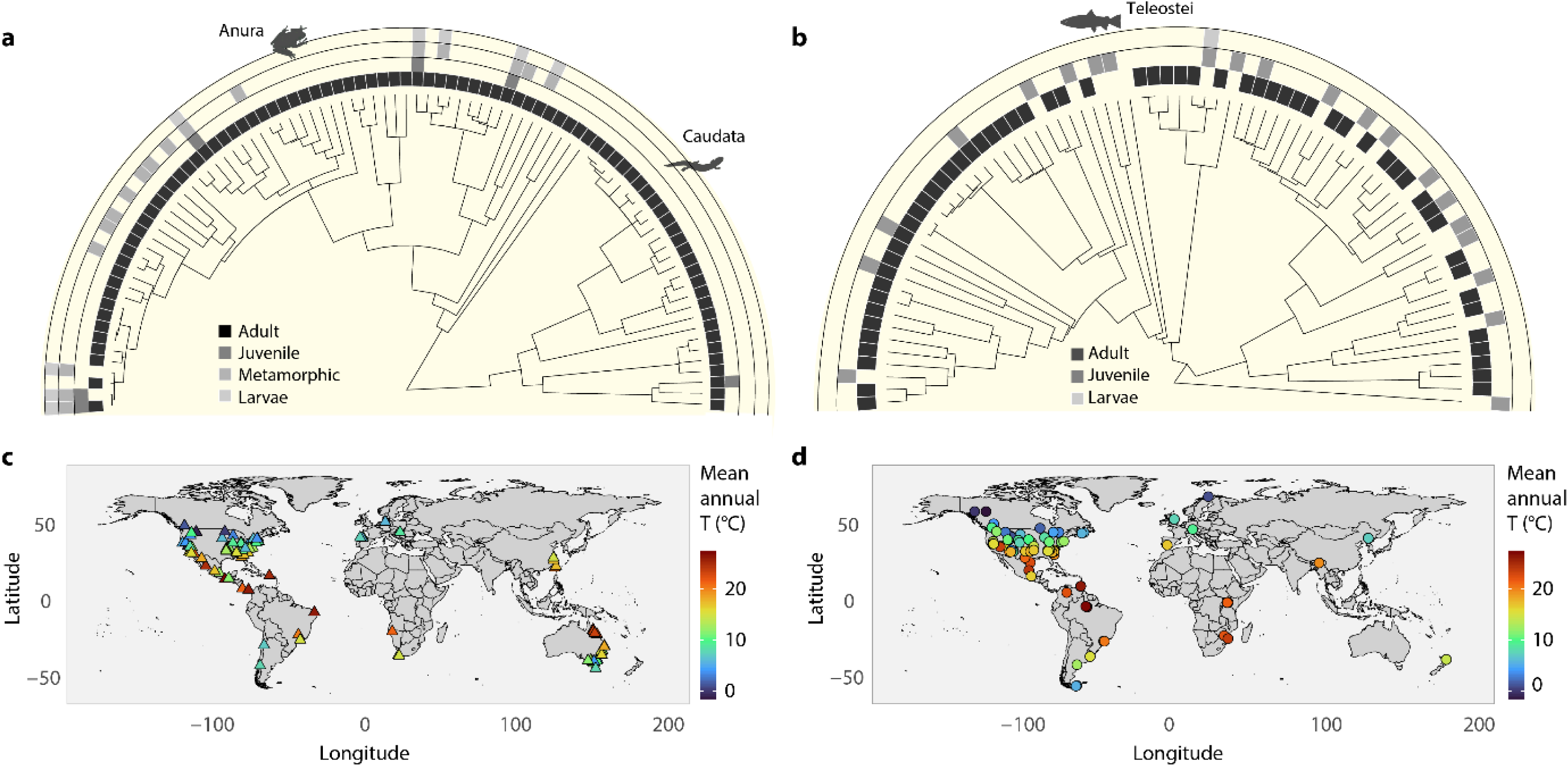
Phylogenetic, ontogenetic, and geographic background of critical thermal maximum (CT_max_) and acclimation response ratio (ARR) measurements in (a, c) amphibians (Anura and Caudata) and (b, d) freshwater fishes (Teleostei). Phylogenetic trees contain (a) 96 amphibian and (b) 93 teleost species assigned to 2 and 14 orders, respectively. Grey shaded squares indicate which life stages of a species were studied. Maps show the geographical origin and local mean temperature of (c) amphibian and (d) fish species. Phylogenetic coverage was approximately 1% within each taxon. Silhouettes were taken from PhyloPic (www.phylopic.org).

Across all species studied, CT_max_ ranged from 20.75 °C to 46.00 °C (Table S1; Fig. S2a). Acclimation capacity in upper thermal tolerance was positive in 97.1% of the experimental studies with an average of 0.22 °C with every 1 °C increase in acclimation temperature across all experimental studies and life stages, whereas the acclimation response for CT_max_ was negative (i.e., negative ARR) in 2.0% of the experimental studies. Across all species studied, ARR ranged from -0.08 to 1.68 (Table S1; Fig. S2b).

For CT_max_ data of freshwater teleosts and amphibians (345 observations = number of experiments), the CIT model (73% explained variance) resulted in a classification tree with 8 internal splits and 9 terminal nodes (Fig. 2). Phylogeny (order/taxon), acclimation temperature and acclimation duration were the main discriminators of CT_max_, followed by geographic origin (annual habitat temperature, bio1) and life stage. The first split separated a group of teleost studies (mainly salmonid species) with lower CT_max_ values compared to the mean of all other studies (p < 0.001). In this group of cold-water fish, CT_max_ increased significantly with acclimation duration (nodes 1-3), with classification thresholds of split 2 and 3 defining differences between short (< 6 days), intermediate (6-21 days) and long acclimation times (> 21 days). For the majority of teleost and amphibian data (308 observations), CT_max_ was significantly dependent on acclimation temperature (split 4, p < 0.001), mean annual habitat temperature (Bio 1; split 5, p = 0.001) and life stage (split 6, p < 0.001). Overall, there was a positive correlation between acclimation/mean annual habitat temperature and CT_max_ (node 4-9). Split 4 separated teleosts and amphibians acclimated below or above 28°C. In the colder group (split 5), CT_max_ was higher for teleosts and amphibians that originated from warmer habitats, with a classification threshold of 16.2°C (node 4 & 5). For species acclimated above 28°C, CT_max_ differed between ontogenetic life stages (split 6), with larvae having higher tolerance limits than juveniles, metamorphs and adults. However, this classification was phylogenetically unbalanced, as all larvae were anuran species (node 8 & 9). Splits 7 and 8 show that after accounting for ontogenetic differences, CT_max_ was higher at warmer acclimation temperature (>31 °C, node 6 & 7) and higher mean annual habitat temperatures (> 16.8 °C, node 8 & 9).

**Figure 2.**
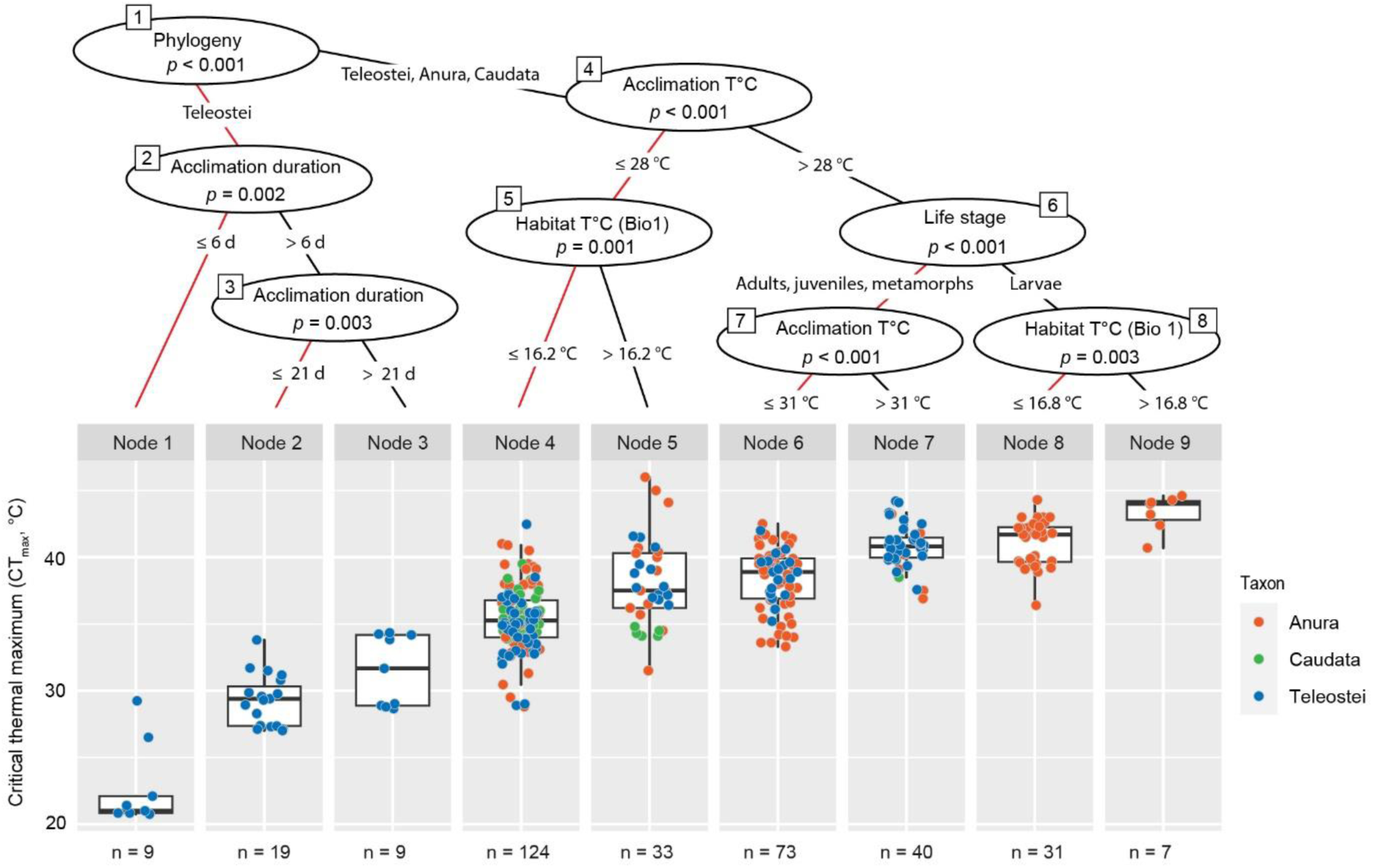
Conditional inference tree (CIT) for critical thermal maximum (CT_max_) of freshwater fish (Teleostei) and amphibians (Anura and Caudata). The sequence of internal nodes (splits) corresponds to a hierarchical structure of significant predictor variables. At each split, lines indicate the classification into groups with higher (black) and lower (red) CT_max_ values. Boxplots show the distribution of CT_max_ values (colored symbols) for each terminal node (n = number of experimental studies). Acclimation T = mean acclimation temperature used in an experiment before CT_max_ measurements (°C). Habitat T = mean annual temperature Bio 1 (°C). Acclimation duration = mean exposure time to acclimation temperatures before CT_max_ measurements (d).

For ARR data (345 observations), the CIT model (52% explained variance) produced a classification tree with 7 internal splits and 8 terminal nodes (Fig. 3). The main discriminators of ARR were phylogeny (order/taxon) and acclimation time, followed by CT_max_ methodology (ramping rate) and a minor influence of local temperature variability (isothermality, Bio 3). The first split of the classification tree separated a large number of teleost studies (102 observations, 68 species, node 1 & 2) with a higher ARR compared to the average of the other fish and amphibian studies in the dataset (p < 0.001). The relatively high thermal plasticity of the fish species in node 1 & 2 was not related to any of the variables examined. Within this group, however, ARR decreased at higher acclimation temperatures, with a classification threshold of 21°C (split 2, p = 0.003). Split 3 was based on CT_max_ methodology, classifying fish and amphibian studies into groups with an acclimation duration longer or shorter than 14 days (node 3-8). Longer acclimation generally resulted in higher ARR values (node 3-5, p < 0.001), but the acclimation effect differed according to ramping rate (split 4), phylogeny (split 5 & 6) and climate variability (split 7). When the acclimation duration exceeded 14 days, ARR studies were further classified according to ramping rate (split 4, p <0.001) and taxon (split 5, p = 0.031). These classifications imply higher ARR values of anurans at slow heating rates (< 0.1°C min^-1^), and higher ARR values of teleosts compared to amphibians at ramping rates > 0.1°C min^-1^ (Fig. 3). If the acclimation period was less than 14 days, a subsequent split occurred according to phylogenetic order (split 6, p < 0.001) and isothermality (split 7), indicating a positive relationship between ARR and local temperature variability (node 7 & 8, p = 0.011). Ontogenetic differences in ARR were not detected.

**Figure 3.**
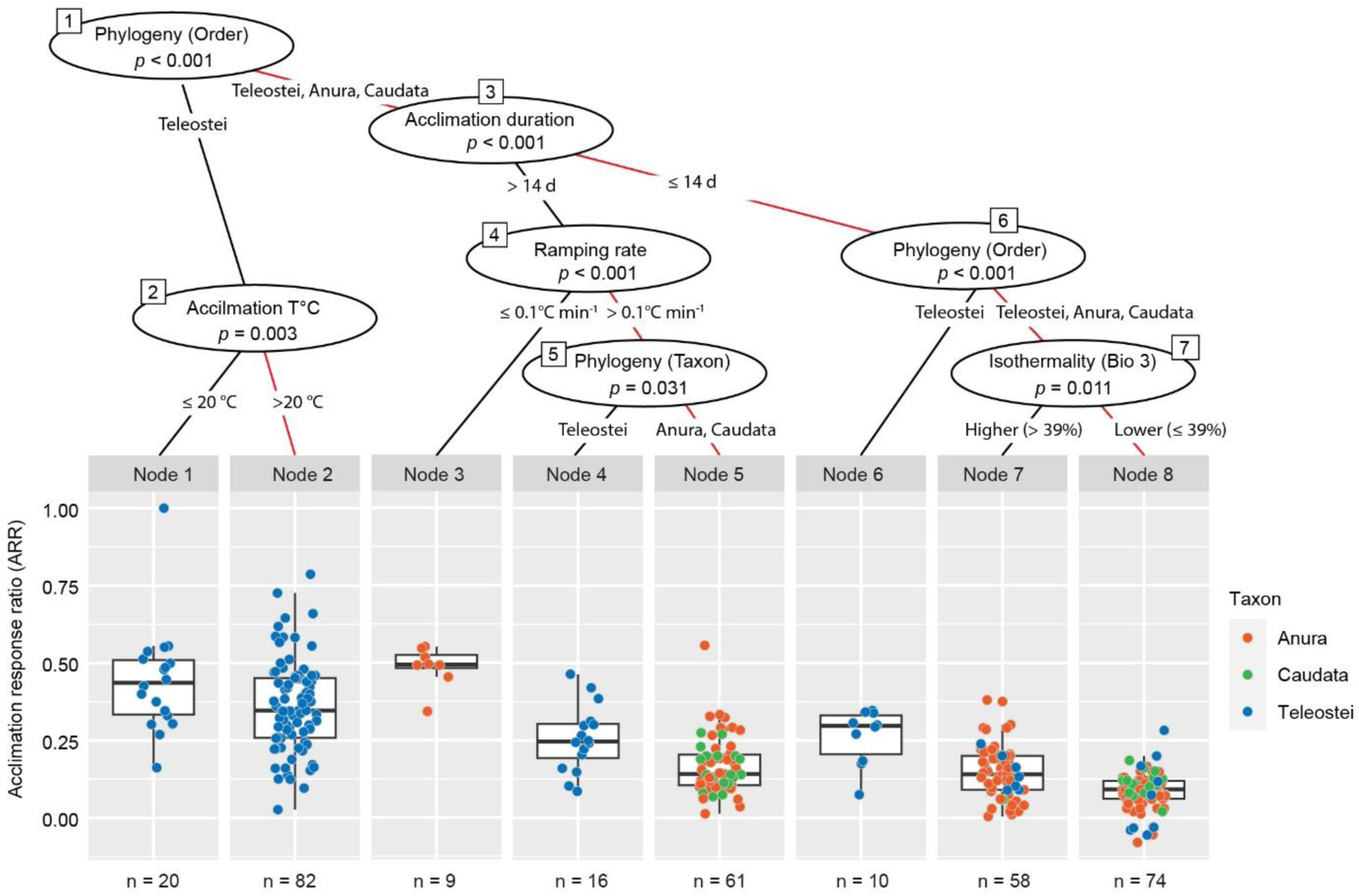
Conditional inference tree (CIT) for the acclimation response ratio (ARR) of freshwater fish (Teleostei) and amphibians (Anura and Caudata). The sequence of internal nodes (splits) corresponds to a hierarchical structure of significant predictor variables. At each internal node, lines indicate the classification into groups with higher (black) and lower (red) ARR values. Boxplots show the distribution of ARR values (colored symbols) for each terminal node (n = number of experimental studies). In node 1, high outlier values (n = 2) were not shown for scaling reasons. Acclimation T = mean acclimation temperature used in an experiment before CT_max_ measurements (°C). Ramping rate = heating rate for CT_max_ measurements in °C per minute. Isothermality = local temperature variability Bio 3 (%). Acclimation duration = mean exposure time to acclimation temperatures before CT_max_ measurements (d).

Isothermality (Bio 3; R² = 0.708, p <0.001) and annual mean temperature (Bio 1; R² = 0.696, p < 0.001) decreased from the tropics to the polar latitudes (Fig. S4). Both bioclimatic variables did not correlate with elevation (Bio 1: R² = 0.137, p <0.001; Bio 3: R² = 0.012, p = 0.046).

## 4. Discussion

### 4.1 Context-Dependent Drivers and Broad-Scale Patterns of Physiological Limits

The most striking result from our analyses is that the acclimation capacity of freshwater fish was more than twice that of amphibians, indicating a strong phylogenetic signal. Our findings align with those reported in previous syntheses (Gunderson & Stillman 2015; Rohr et al. 2018; Morley et al. 2019; Pottier et al. 2022a), demonstrating a higher thermal plasticity in organisms inhabiting aquatic habitats compared to their terrestrial counterparts in general, and with fish (marine and freshwater) exhibiting greater thermal plasticity compared to amphibians (Gunderson & Stillman 2015). Compared to most (post-metamorphic) amphibians, freshwater fish are restricted to their aquatic habitat throughout their life cycle (Comte & Grenouillet 2013). As aquatic habitats tend to have less spatial variability in operative thermal conditions than terrestrial habitats (Gunderson & Stillman 2015; Sunday et al. 2014), behavioral thermoregulation is constrained, and freshwater fish are more likely than amphibians to use thermal plasticity to buffer against changing thermal conditions. Consequently, the capacity for thermal plasticity appears to be phylogenetically conserved between both taxa (Angilletta et al. 2002; Bodensteiner et al. 2021), depending on the ability for behavioral thermoregulation as explained by the *Bogert effect* (Bogert 1949; Cowles & Bogert 1944; Muñoz 2022), rather than on the level of thermal variation to which a population is exposed to (Huey et al. 1999) as suggested by the *climate variability hypothesis* (Janzen 1967). Moreover, unlike freshwater fishes, most amphibians undergo a habitat shift from a larval aquatic to a post-metamorphic terrestrial habitat (Shi 2000). As an adaptive response to often shallow or temporary larval habitats (Newman 1992), amphibian larvae display a high degree of plasticity in growth and development (Kulkarni et al. 2017; Ruthsatz et al. 2018a,b; Burraco et al. 2021; Sinai et al. 2022), providing a means for increasing fitness (Schlichting & Pigliucci 1998). Therefore, plasticity in timing of metamorphosis appears to be more important than that in thermal tolerance to reduce mortality risk (Rudolf & Rödel 2007) due to desiccation or temperature extremes (Burraco et al. 2022; Albecker et al. 2023). This might, at least in part, explain the lack of an ontogenetic pattern in acclimation capacity in amphibians.

Unlike acclimation capacity, geographic origin was one of the primary determinants of heat tolerance in both taxa, with a higher heat tolerance in populations from regions with a higher annual mean temperature compared to populations from regions with a lower annual mean temperature. These findings support, at least in part, the *climate variability hypothesis* (Janzen 1967; Bozinovic et al. 2011) since annual mean temperature decreases with latitude and is highest in tropical low-elevation regions. Both taxa, therefore, exhibit physiological adaptations to latitude-dependent thermal regimes to which they are exposed. Our results agree with the findings of a large body of research that has confirmed the link between physiological limits and large-scale geography based on a species’ local adaptation to temperature and other associated climatic variables (e.g., Gutiérrez-Pesquera et al. 2016; Sunday et al. 2011, 2019; Pintanel et al. 2022; but not: Addo-Bediako et al. 2000; Sørensen et al. 2016). Yet, in addition to the broader literature, our synthesis indicated that CT_max_ was not only *generally* higher in populations from warmer regions as often found in low latitudes but also, importantly, varied with ontogeny. These findings emphasize the importance of assessing life stage-specific sensitivity to thermal stress as well as spatial climatic differences in conservation science. Therefore, focusing on large-scale geographical patterns for predicting how biodiversity will respond to future environmental change might bear the risk of overlooking context-dependent variation in thermal traits and thus, intraspecific differences in vulnerability to changing thermal conditions (Duarte et al 2012, Gutierrez-Pesquera et al 2016). For example, Bovo et al. (2023) demonstrated that responses of tropical amphibians to climate variation were heterogenous as a consequence of intraspecific variation in physiological traits and spatial variation in climate with elevation. Furthermore, Sunday et al. (2011) and Pinsky et al. (2019) reported that the physiological sensitivity of ectotherms across all latitudes depended on the realm, with terrestrial ectotherms being less sensitive to warming than aquatic ectotherms due to their higher capacity for behavioral thermoregulation.

### 4.2 Life Stage-Specific Thermal Sensitivity as a Key Factor in Species Vulnerability to Climate Change

In species with complex life-histories such as amphibians and teleost fish, life stages differ in size, morphology, physiology, and behavior (Wilbur 1980). Therefore, selection might promote stage-specific adaptations in thermal physiology (Enriquez-Urzelai et al. 2019; Ruthsatz et al. 2022a -or b). Ignoring those life stage-specific differences in thermal physiology may drastically underestimate climate vulnerability of species with consequences for successful conservation actions. Here, we found that CT_max_ but not acclimation capacity differed between life stages in amphibians, with a higher CT_max_ in larval stages. However, this pattern was only evident when high acclimation temperatures were used in experimental set-ups and was most pronounced in larvae from habitats with a high annual mean temperature as those found in the tropics. Limnic larvae may have a reduced capacity for behavioral thermoregulation due to their limited body size impairing the movement between different microclimates (Kingsolver et al. 2011; Sinclair et al. 2016; Enriquez-Urzelai et al. 2019), making them more dependent on passive responses to temperature fluctuations. To cope with changes in temperatures, a high heat tolerance is therefore advantageous in early life stages (Ruthsatz et al. 2022a). This might be particularly true for animals that are already living close to their heat tolerance since the ontogenetic difference was only evident at high acclimation temperatures and strongest in anurans from habitats with a high annual mean temperature. In contrast, post-metamorphic stages might rather be able to select favorable microclimates by behavioral thermoregulation (Navas et al. 2007; Haesemeyer 2020). This is particularly true for amphibians, as their post-metamorphic terrestrial habitats offer much spatial variability in operative thermal conditions (Gunderson & Stillman 2015), while juvenile and adult (freshwater) fish are able to behaviorally thermoregulate by performing vertical and horizontal movements (Amat-Trigo et al. 2023; Breau et al. 2007; but not: Clark et al. 2022). Our findings are in line with the pattern found for aquatic larvae by Cupp (1980), Sherman (1980), Enriquez-Urzelai et al. (2019), and Ruthsatz et al. (2022a), who demonstrated a higher CT_max_ in amphibian larvae than in post-metamorphic stages. In contrast, Dahlke et al. (2020) found no difference in heat tolerance between larval and adult stages in marine and freshwater fish. Notably, our synthesis yielded only one estimate for larval CT_max_ and acclimation capacity in freshwater fish and, thus, we lack the data for any conclusion. Given that small body sizes of larvae restrict their capacity for behavioral thermoregulation, we would would expect freshwater fish to exhibit the same life stage-specific differences in thermal sensitivity observed in amphibians. In a recent study on the European common frog (*Rana temporaria*), young larvae may define the climate sensitivity of populations since that life stage exhibited the lowest acclimation capacity (Ruthsatz et al. 2022a). Furthermore, it is worth noting that our synthesis did neither encompass embryos nor gametes producing/spawning stages, which have recently been reported to have the lowest heat tolerance in fish (Dahlke et al. 2020, 2022; Pottier et al. 2022b) and the lowest acclimation capacity across ectotherms (Pottier et al. 2022a). To better identify potential life history bottlenecks in thermal sensitivity in amphibian, fish and other taxa inhabiting freshwater, future studies should adopt a more comprehensive approach by considering a wider range of life stages within species.

### 4.3 Understanding Context-Dependent Physiological Adaptation in Ectotherms

Global syntheses on physiological studies can help us determine the winners and losers of climate change through assessment of broad-scale patterns of species’ thermal limits and acclimation capacity for modifying their thermal tolerance (Somero 2010). This knowledge, in turn, enables us to develop suitable conservation strategies to mitigate the negative effects of climate change. However, our key findings emphasize that assessing species’ vulnerability to changing thermal conditions based on large-scale geographic and/or phylogenetic patterns in thermal traits might cover up context-dependent physiological adaptations. In other words, tropical ectothermic species are considered particularly vulnerable to global warming as they live close to their physiological limits and have poor acclimation ability (Tewksbury et al. 2008; Huey et al. 2009; Sunday et al. 2014), but such generalizations might for instance ignore intraspecific variation in physiological limits across variation in climate with elevation (Bovo et al. 2023). Physiological adaptations are driven by the interplay between microclimate temperature heterogeneity and the behavioral thermoregulatory abilities of ectotherms (Huey et al. 2012; Pincebourde et al. 2016; present study) depending on their habitat characteristics (Pinsky et al. 2019; Kulkarni et al. 2017), ontogeny (Enriquez-Urzelai et al. 2019; Ruthsatz et al. 2022a), life history traits such as body size (Rubalcaba et al. 2020; Peralta-Maraver & Rezende 2021) or activity patterns (Navas et al. 2007; Ruthsatz et al. 2022b), and/or energy balance (Pörtner et al. 2005; Muñoz et al. 2022). Physiological traits and limits are consequently rather evolutionary driven dynamic concepts than fixed values for a species (Bovo et al. 2018, 2023; Navas et al. 2022). In order to improve predictions of climate change impacts on biodiversity, it is imperative to deepen our understanding of context-dependent physiological adaptations (Meek et al. 2023), thereby advancing the development of suitable conservation measures/strategies that incorporate evolutionarily enlightened perspectives (Ashley et al. 2003; Cook & Sgró 2018) beyond the species level (Fig. 4).

**Figure 4.**
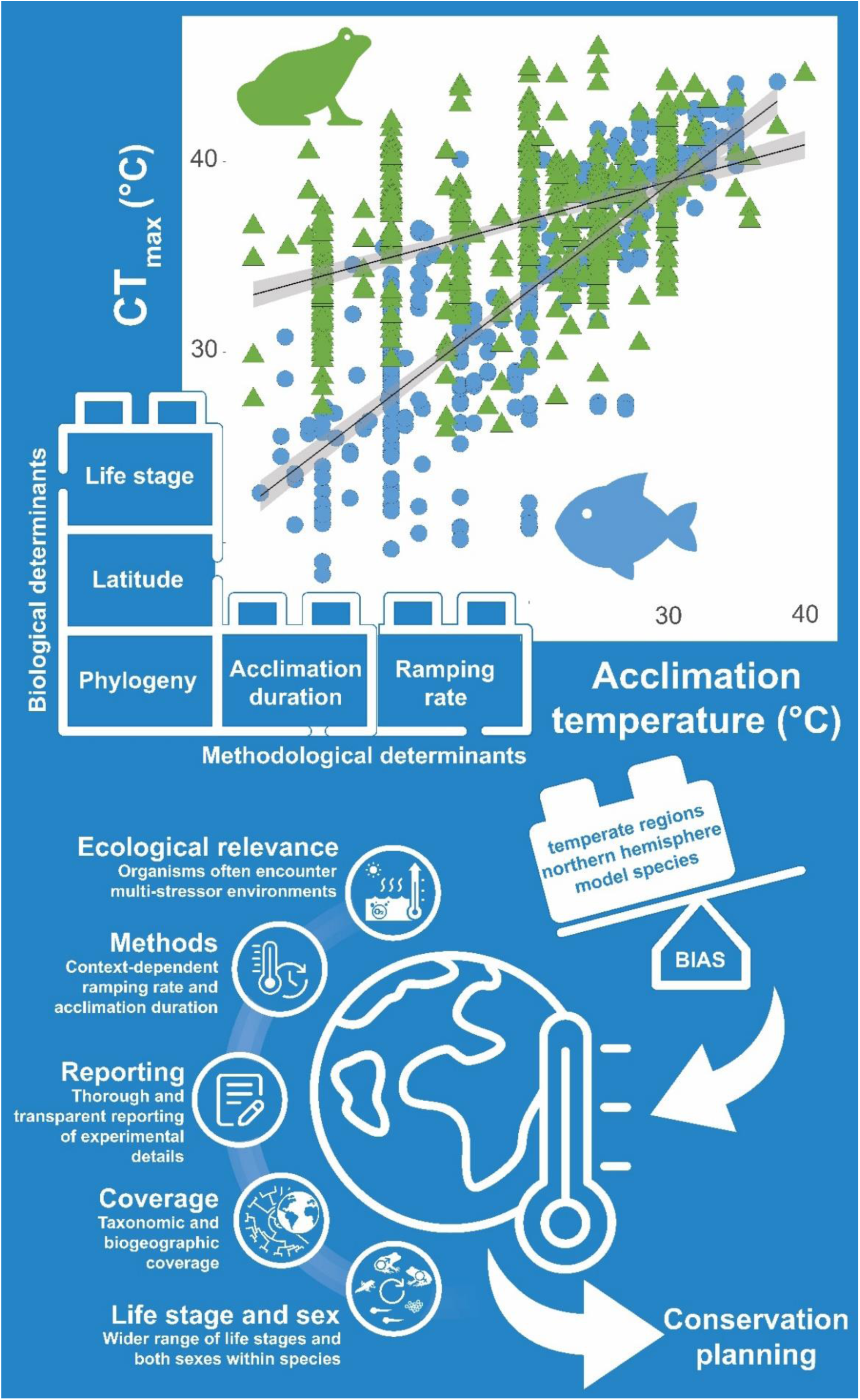
Synthesis of biological and methodological determinants as well as key directions for future research on acclimation capacity in amphibians and freshwater fish to advance the application of thermal traits in assessing species’ and populations’ vulnerability to climate change. See text for further details.

### 4.4 Conclusion

There is a growing body of physiological studies assessing thermal limits and acclimation capacity of species, investigating physiological systems setting these limits to better predict shifts in the productivity and species distribution patterns in a warming world. Our synthesis points to representation biases in taxonomy, species’ biogeographic distribution, life stage, and biases resulting from non-standardized study design. Here, we found the influence of life stage, phylogeny, and thermal adaptation to depend on acclimation duration and acclimation temperature of the animals and the ramping rate used, underscoring the importance of a thoughtful selection of the methodological approach. Notably, due to the biases present in existing data sets, most results from subsequent analyses (including those presented here) cannot be generalized to all geographic regions, life stages, and (taxonomic) groups, which limits the ability to predict organismal responses to global change (White et al. 2021). Therefore, we conclude our synthesis by addressing those data inadequacies and proposing methods to enhance data collection presented in five themes (Fig. 4):

#### Biogeographic and taxonomic coverage

We found geographical trends in physiological limits associated with bioclimatic conditions at different latitudes. Thus, additional research is needed in poorly represented (mostly tropical) regions that generally correspond to low- and middle-income countries (King 2004). Addressing this issue can yield additional, important insights, as such geographic bias in existing data sets has been shown to limit our ability to predict organismal responses in these less-studied regions (White et al. 2021). Our synthesis highlights that most experiments have been conducted in North America, Europe, and Australia and information gaps exist for most parts of Neotropics, Africa and Sino-Oriental regions. Those regions harbor the highest diversity of species for both anurans and freshwater fishes (Jenkins et al. 2013, Van der Sleen & Albert, 2022). Such regional differences in research effort are common in conservation science (Schiesari et al. 2007; Winter et al. 2016; White et al. 2021; McLaughlin et al. 2022; Sinai et al. 2022) despite the fact that under-studied regions contain the vast majority of global biodiversity hotspots (Mittermeier et al. 2011). Furthermore, most of these experiments used species that are common, widely distributed, and/or easily obtained by researchers. Studies on other species (particularly those already in decline) are needed to avoid taxonomic bias and reach stronger conclusions on whether specific taxa might be more sensitive to global warming (da Silva et al. 2020). Additionally, there may be an elevation bias in sampling, as most available data represent lowland rather than mountain species. As, for example, CT_max_ of tropical and subtropical amphibian species has been shown to vary within the same latitude (Duarte et al 2012, Gutierrez-Pesquera et al 2016), we urge future studies to focus on elevation gradients as well as cross-latitudinal sampling, especially in tropical high-mountain populations (Bovo et al., 2023).

#### Relevance of life stage and sex

We urge future studies to measure a wider range of life stages and to measure both sexes within species to better identify thermal bottlenecks and assess climate change risks (Klockmann et al. 2017; Dahlke et al. 2020). All of the summaries to date likely overestimate physiological limits since most studies have been performed on adults, and thermal tolerance increases (Klockmann et al. 2017; Rubalcaba & Olalla-Tárraga 2020; but not: di Santo & Lobel 2017) and acclimation capacity decreases (Pottier et al. 2021) with body size. Furthermore, after sexual maturity, females and males differ in a wide range of morphological, physiological, and behavioral aspects as well as in their energetic investment in gamete production (Hayward & Gillooly 2011). A recent meta-analysis across ectothermic taxa revealed that the acclimation capacity differed between adult males and females in wild-caught animals (Pottier et al. 2021). Furthermore, Dahlke et al. (2020) found narrower thermal tolerance ranges in spawning females and van Heerwaarden and Sgrò (2021) demonstrated that a low heat tolerance of male fertility is a critical bottleneck in insects.

#### Ecological relevance

Future laboratory experiments should adopt a more comprehensive and multifaceted approach for higher ecological relevance of thermal trait estimates (Desforges et al. 2023). Under natural conditions, organisms must often cope with multiple simultaneously occurring environmental stressors (Rohr & Palmer 2013; Gunderson et al. 2016) such as declining dissolved oxygen levels in freshwater habitats due to climate-induced temperature increases (Pörtner & Peck 2010). As thermal limits are shaped by oxygen availability (Pörtner 2001, 2010), organisms might exhibit lower thermal limits under natural conditions. Moreover, exposure to pollutants might reduce thermal tolerance (Little & Seebacher 2015) or acclimation capacity (Ruthsatz et al. 2018b) due to increased metabolic demands of detoxification processes or disruption of endocrine pathways involved in physiological acclimation. Physiological responses to increased water temperature as performed in the experimental studies summarized here (i.e., a single stressor) may not align with observed responses of individuals in the natural environments with multiple stressors (Katzenberger et al. 2014; Potts et al. 2021).

#### Methodological approach

Customizing protocols to account for organismal and context-dependent variations in physiological limits (e.g., body size, life stage, sex, thermal history) will allow researchers to obtain more ecologically relevant estimates to inform conservation efforts. The application of an acute thermal ramping rate and a standardized endpoint such as the loss of equilibrium are used to measure critical thermal limits (Becker & Genoway 1979). The estimates are sensitive to differences in the methods. For example, faster ramping (heating) rates tend to yield higher thermal tolerance estimates compared to slower ramping rates (Moyano et al. 2017; Kovacevic et al. 2019; Penham et al. 2023). Furthermore, measuring several endpoints (loss of equilibrium/rightening response, onset of spasms/heat rigor) if possible (Wu & Kam 2005; Cowan et al. 2023) might facilitate comparisons between different taxonomic groups and/or experiments (Lutterschmidt & Hutchison 1997). Using wild-collected animals is important as those reared in the laboratory may have physiology traits that differ from wild conspecifics (Pottier et al. 2021; Morgan et al. 2022). Methodological recommendations have been recently published and comparable methods are needed to compare thermal limits of different life stages (Cowan et al. 2023; Desforges et al. 2023).

#### Through reporting of research details

Thorough and transparent reporting of experimental details in empirical studies such as sampling location and animal origin, among others, is required to enhance the comparability of studies on thermal traits. Furthermore, most studies working on adults did not report the sex of animals despite the potential for sex (or reproductive state) to be important factors in thermal sensitivity. Additionally, reporting microhabitat temperatures (i.e., in-site temperatures) will improve our understanding of thermal adaptation at the population-level. The ability to make broad-scale comparisons of thermal tolerance across taxa, life stages, and regions will be enhanced when studies report as much methodological detail as possible. Consequently, these future studies will contribute more robust estimates of climate vulnerability needed to guide climate change interventions.

By considering our recommendations, future studies will be more comparable, facilitating the utilization of respective findings in large-scale studies and models that assess species vulnerability and thus, population dynamics under global warming.

## 5. Data availability

Data associated with this study are available in the figshare data repository under https://figshare.com/s/57775032a0b79c416ef2 (DOI: 10.6084/m9.figshare.24872133; published after acceptance).

## 7. Acknowledgements

We would like to thank the German Research Foundation (DFG) for providing financial support to KR and PCE for this study and the Universität Hamburg for providing financial support to KR by the PostDoc1st award. We thank M. Multsch for assisting with the data visualization and U. Enriquez-Urzelai and M. H. Bernal for providing data on request. We thank Trina Rytwinski for her guidance on the statistical approach. We are grateful for the valuable comments provided by two anonymous reviewers. We are particularly grateful to all authors of the articles included in our systematic literature review, whose effort and work made the present study possible.

## 8. Author contributions

Conceptualization: KR and MAP. Methodology: KR, KA, FD, SW, SG, MAP. Data Extraction and Quality Check: KR, KA, PCE, MLL, FD and SW. Formal Analysis: FD and SW. Investigation: KR, FD, and SW. Data Curation: KR. Visualization: KR, FD, and SW. Writing – Original Draft: KR, FD, and MAP. Writing – Review and Editing: all authors. Funding Acquisition: KR. Project Administration: KR. Supervision: KR and MAP. All authors gave their final approval for submission.

## 10. Statement of Ethics

The authors have no ethical conflicts to disclose.

