## Supplementary Material for "Acclimation Capacity to Global Warming of Amphibians and Freshwater Fishes: Drivers, Patterns, and Data Limitations"

**Global Change Biology – Revised Manuscript**

***Running title:*** *Acclimation: Amphibians and Freshwater Fishes*

Katharina Ruthsatz[1,2], Flemming Dahlke[3], Katharina Alter[4], Sylke Wohlrab[5,6], Paula C. Eterovick[1], Mariana L. Lyra[7,8], Sven Gippner[1], Steven J. Cooke[9], Myron A. Peck[4,10]

*[1]Zoological Institute, Technische Universität Braunschweig, Mendelssohnstraße 4, 38106 Braunschweig, Germany*

*[2]Institute of Zoology, Universität Hamburg, Martin-Luther-King-Platz 3, 20146 Hamburg, Germany*

*[3]Ecology of Living Marine Resources, Universität Hamburg, Große Elbstraße 133, 22767 Hamburg, Germany*

*[4]Department of Coastal Systems, Royal Netherlands Institute for Sea Research, PO Box 59 1790 AB, Den Burg (Texel), the Netherlands*

*[5]Alfred Wegner Institute Helmholtz Center for Polar and Marine Research, 27570 Bremerhaven, Germany*

*[6]Helmholtz Institute for Functional Marine Biodiversity at the University of Oldenburg (HIFMB), 23129 Oldenburg, Germany*

*[7]New York University Abu Dhabi, Saadiyat Island, Abu Dhabi, P.O. Box 129188, United Arab Emirates[8]Center for Research on Biodiversity Dynamics and Climate Change, State University of São Paulo-UNESP, Rio Claro, SP, Brazil.*

[9]*Fish Ecology and Conservation Physiology Laboratory, Department of Biology and Institute of Environmental and Interdisciplinary Science, Carleton University, Ottawa, ON K1S 5B6, Canada*

[10]*Marine Animal Ecology Group, Department of Animal Sciences, Wageningen University, PO Box 338, 6700 AH, Wageningen, the Netherlands*

---

Key words: *acclimation response ratio, $CT_{max}$, thermal tolerance plasticity, developmental phenotypic plasticity, thermal bottleneck, metamorphosis, Bogert Effect, climate variability hypothesis*

Corresponding author: **Katharina Ruthsatz**; ORCID: 0000-0002-3273-2826. Current address: Zoological Institute, Technische Universität Braunschweig, Mendelssohnstraße 4, 38106 Braunschweig, Germany. Phone: 0049 531 3912393..

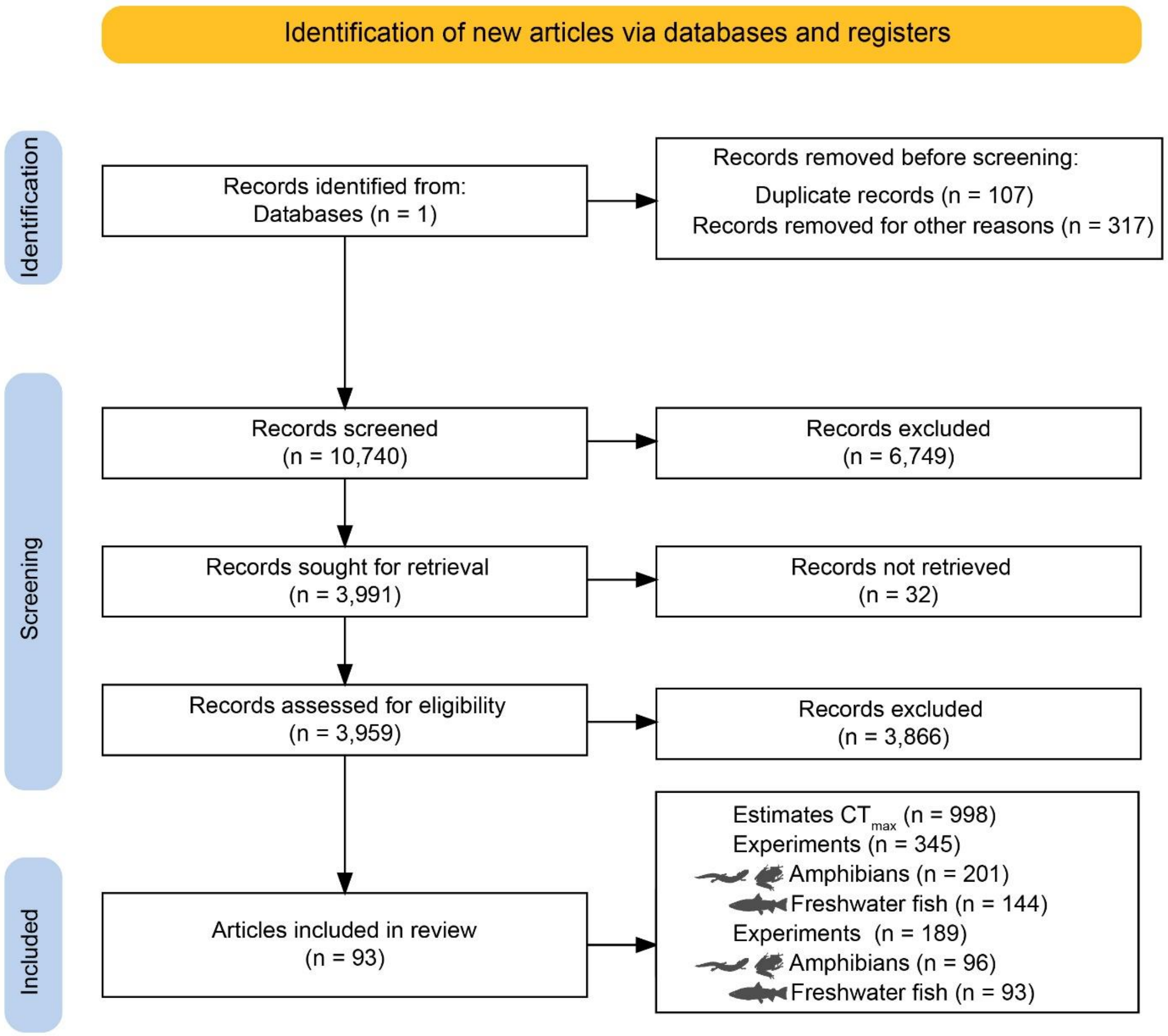


**Figure S1**. PRISMA flow diagram adapted from Page et al. (2021) showing literature search procedures and screening processes (created with Shiny app, Haddaway et al. 2022).

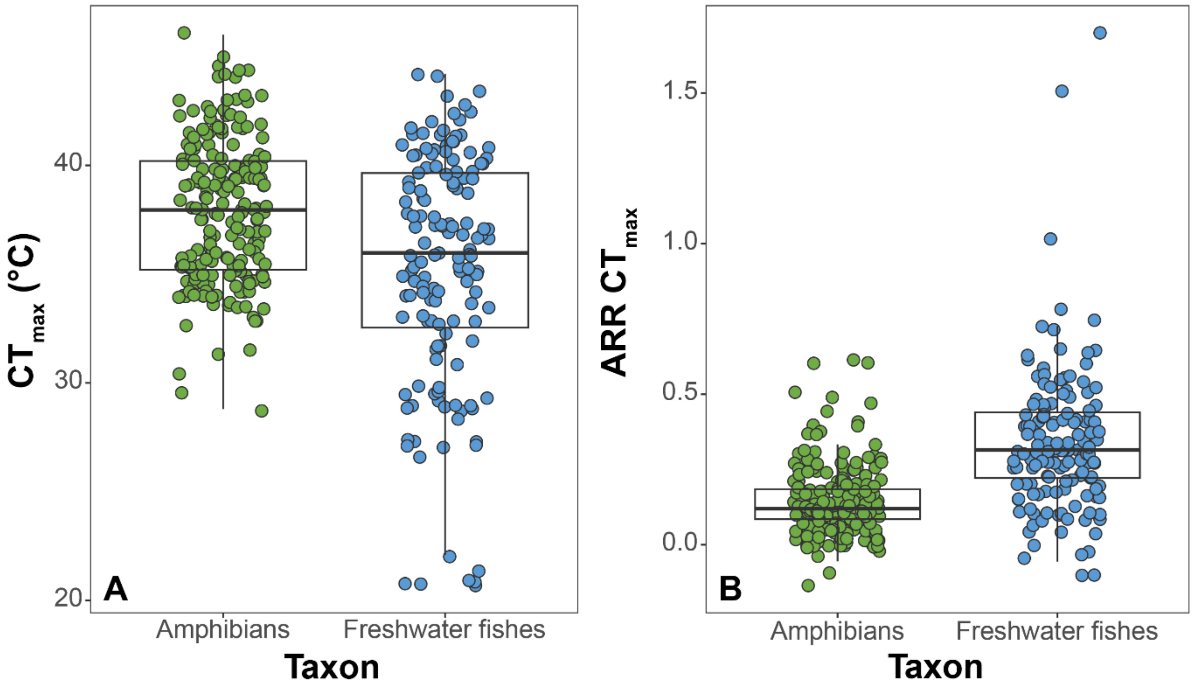


**Figure S2.** Taxon-specific **A** critical thermal maximum ($CT_{max}$) and **B** Acclimation response ratio of $CT_{max}$ (ARR $CT_{max}$) for amphibians and freshwater fishes. Green: Amphibians. Blue: Freshwater fishes. Numbers in parentheses = sample size.

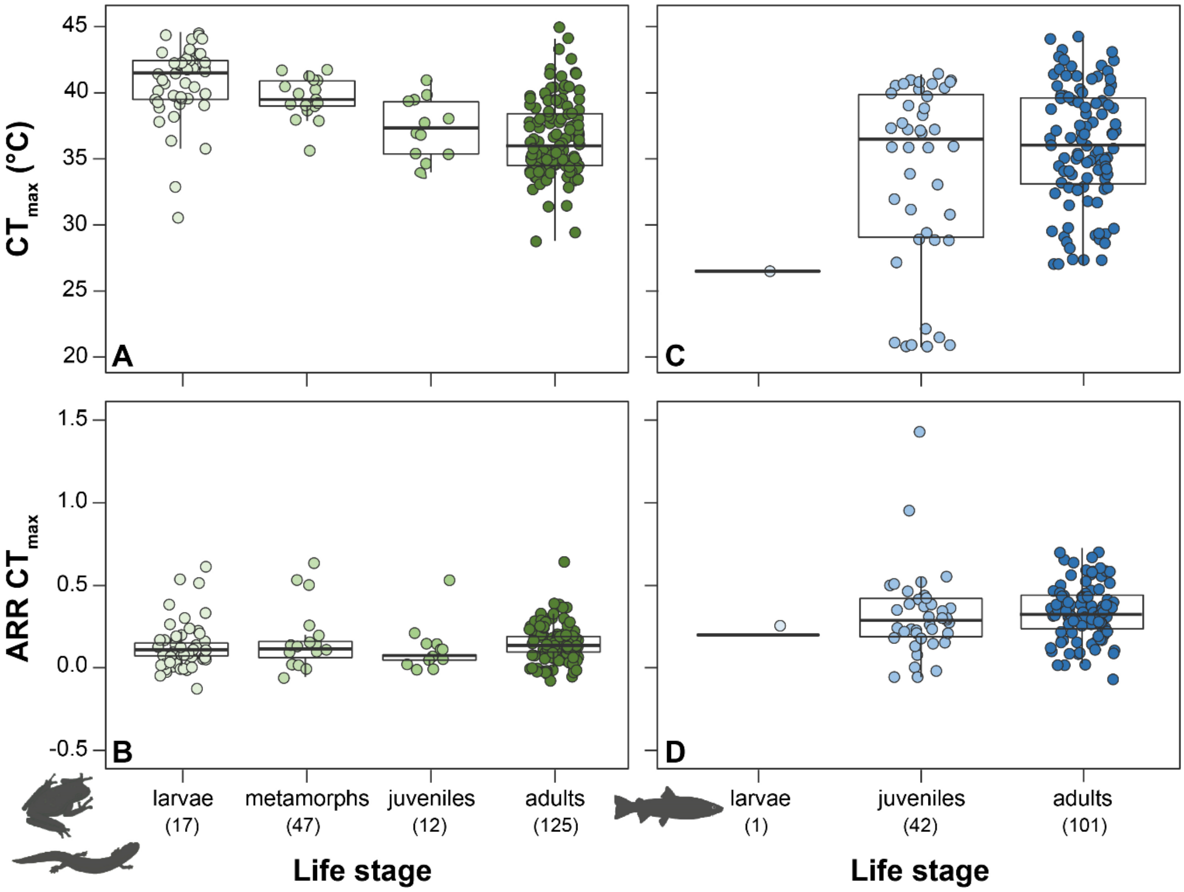


**Figure S3. AC** Life stage-specific critical thermal maximum ($CT_{max}$) and **BD** Acclimation response ratio of $CT_{max}$ (ARR $CT_{max}$) for amphibians and freshwater fishes. Green: Amphibians. Blue: Freshwater fishes. Numbers in parentheses = sample size.

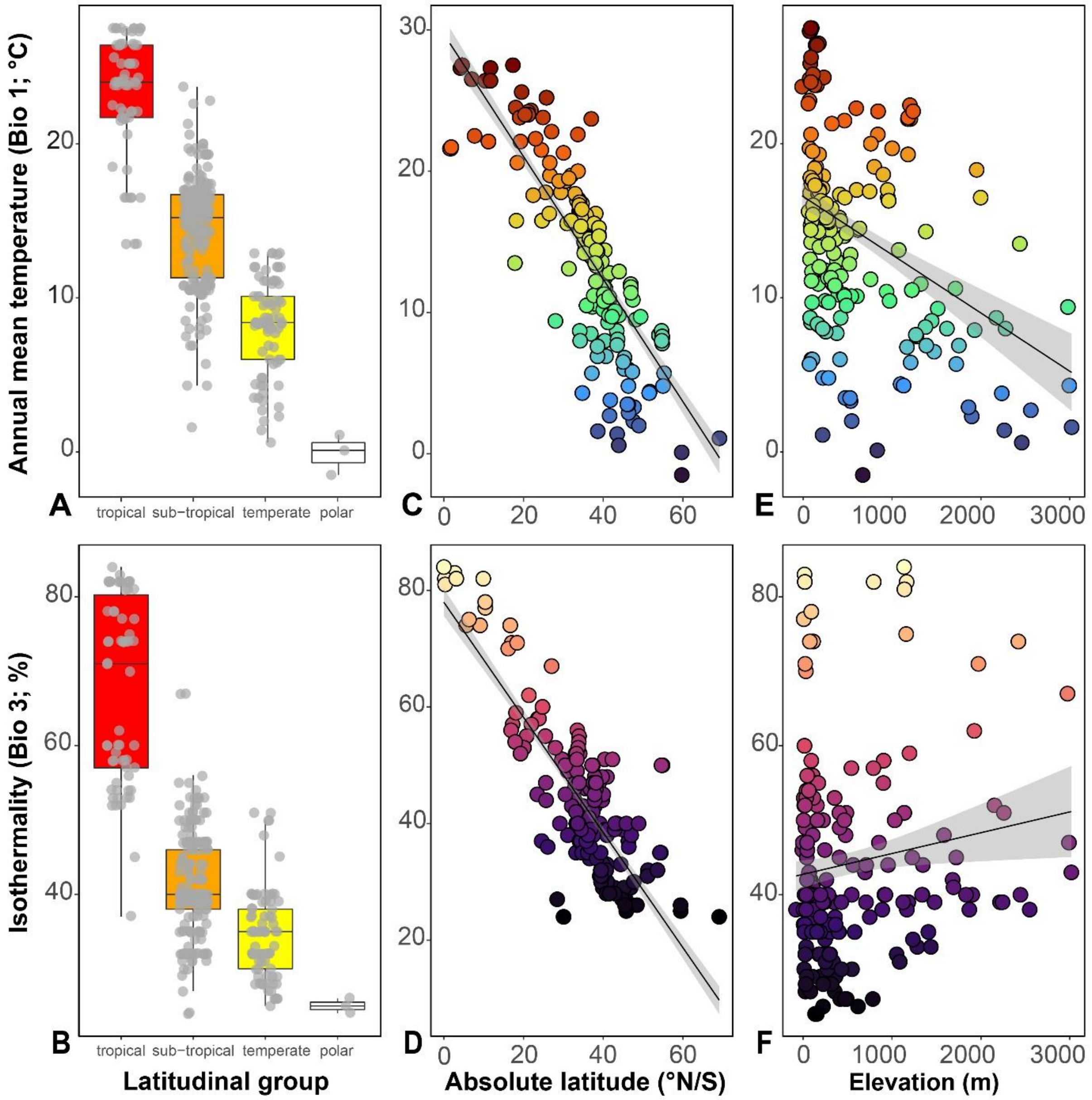


**Figure S4. AB**. Isothermality (Bio 3, %), and **B**. annual mean temperature (Bio 1, °C) range for four latitudinal groups (i.e., tropical, sub-tropical, temperate, and polar) and **CD** as a function of absolute latitude (°N/S) and **EF** altitude (m).
